# ppLM-CO:Pre-trained Protein Language Model for Codon Optimization

**DOI:** 10.1101/2024.12.12.628267

**Authors:** Shashank Pathak, Guohui Lin

## Abstract

Messenger ribonucleic acid (mRNA) vaccines represent a major advancement in synthetic biology, yet their efficacy remains limited by how efficiently the encoded protein is translated within the host. Since multiple codons can code for the same amino acid, the search space of possible coding sequences (CDS) within mRNA grows exponentially with protein length, making the problem highly underdetermined. Finding CDSs that yield efficient translation hinges on codon optimization—the process of choosing among synonymous codons, while encoding the same protein but differ in their effects on translation speed, tRNA availability, and mRNA secondary structure. Recent deep learning approaches have framed codon optimization as a sequence learning problem, where the goal is to model context-dependent codon usage patterns across the amino acid in protein sequence. However, these methods rely on large sequence models that learn amino-acid embeddings from scratch, leading to computationally intensive training. We propose ppLM-CO, a lightweight codon optimization framework that integrates pretrained protein language models (ppLMs) to directly provide contextual amino-acid embeddings, thereby eliminating the need for embedding learning. This design reduces trainable parameters by over 92% – 99% compared with prior deep models while maintaining complete biological fidelity. In-silico evaluations across three species and two vaccine targets—SARS-CoV-2 spike and Varicella-Zoster Virus (VZV) gE viral proteins—demonstrate that ppLM-CO consistently achieves higher expression and competitive stability, establishing a scalable and biologically consistent approach for codon optimization.

## 1 Introduction

Messenger RNA (mRNA) vaccines have emerged as a transformative technology in synthetic biology. For instance, vaccines from Pfizer (BNT162b2) and Moderna (mRNA-1273) proved highly effective in limiting the spread of COVID-19 [1, 2]. Their success has catalyzed the development of mRNA vaccines for diverse pathogens, including varicella-zoster virus (VZV), cancer, malaria, and influenza [3]. However, the efficacy of an mRNA vaccine depends critically on how efficiently the host translates the delivered mRNA into the corresponding antigenic protein, which determines the strength and duration of the immune response. This translation efficiency is governed by two key properties: protein expression, reflecting how effectively ribosomes synthesize the antigen, and mRNA stability, which dictates how long the transcript remains available for translation. Thus, achieving high protein expression and mRNA stability is central to effective vaccine design and therapeutic protein production [4, 5, 6].

However, designing an mRNA sequence remains inherently challenging due to genetic code degeneracy. Each protein comprises amino-acids, and each amino-acid can be encoded by one or more codons—triplets of nucleotides forming the CDS within the mRNA, making it an ill-posed problem. Although different CDSs may translate to the same protein sequence, they differ in codon usage bias, tRNA abundance, and mRNA secondary structure, leading to large variations in expression and stability [7, 8]. Identifying the optimal CDS that maximizes expression while maintaining stability—known as codon optimization [9]—remains a core yet computationally difficult problem due to the combinatorial size of the synonymous CDS space.

Sequential deep learning models like recurrent neural networks (RNNs) and transformers [10], due to their contextual representation capabilities, have been utilized for modelling biological sequences [11]. Recent deep learning approaches address codon optimization by modeling codon selection for each amino-acid as a context-dependent process [12, 13, 14, 15, 16]. These methods either formulate the task as sequence tagging—predicting a codon label for each amino-acid—or as sequence-to-sequence translation, where the amino-acid sequence is treated as a source language and the nucleotide sequence as a target language. In both paradigms, the quality of amino-acid embeddings is critical for capturing contextual dependencies that influence codon choice. However, learning such embeddings from scratch is computationally expensive and we find it to be redundant, given the availability of large-scale pretrained protein language models (pLMs) [17, 18] that already encode rich evolutionary and structural information. Despite their potential, these pretrained representations remain underexplored for codon optimization.

To this end, we propose ppLM-CO—a **P**retrained **P**rotein **L**anguage **M**odel for **C**odon **O**ptimization. It leverages ProtBert [17] as a pretrained protein language model (ppLM) to generate contextual amino-acid embeddings, thus eliminating the need to learn them from scratch. On top of these fixed embeddings, we introduce a biologically constrained generalized linear model (GLM) classifier, which is the only trainable component in our framework. A biological masking as a prior is applied at the output layer to suppress logits of non-synonymous codons, ensuring that predictions remain genetically valid by construction. This formulation keeps intact the sequence-tagging paradigm [12], while simplifying codon optimization into a biologically consistent, lightweight multi-class classification problem. Owing to the lightweight design, ppLM-CO reduces trainable parameters by over 92% – 99% compared to transformer-based methods [16], while still achieving state-of-the-art codon adaptation index (CAI) [19] and competitive minimum free energy (MFE)—proxies for expression and stability [20] respectively—across three host species: Human, Escherichia coli (*E. coli*), and Chinese-Hamster. Furthermore,in *in-silico* mRNA vaccine design for the SARS-CoV-2 spike and VZV gE viral proteins, ppLM-CO generates optimized CDSs that surpasses existing computational methods [16, 21] and publicly available industry vaccines CDSs coming from Pfizer (BNT162b2) and Moderna (mRNA-1273) [22]. These results establish ppLM-CO as an efficient, biologically grounded framework for codon optimization and a promising computational tool for mRNA vaccine and synthetic gene design.

### Availability

ppLM-CO is released as an open-source tool with an integrated Gradio-based graphical interface for interactive use. The tool is available here: https://github.com/shashankcuber/Pre-trained-PPLM-Codon-Optimization

## 2 Material and Methods

### 2.1 Method Overview

In codon optimization, the goal is to generate the CDS (*Y*) for a given protein sequence (*X*). Formally, the dataset 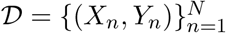, consists of *N* pairs of input protein sequences (*X*_*n*_ = (*a*_1_, …, *a*_*L*_)) and their corresponding CDSs (*Y*_*n*_ = (*c*_1_, …, *c*_*L*_)), where each amino-acid *a*_*i*_ ∈ 𝒜 (20 amino-acids) and each codon *c*_*i*_ ∈ 𝒞 (61 codons). Three stop codons were excluded from the label space since they do not encode amino-acids and are not subject to synonymous variation within the CDS. The objective is to learn the mapping

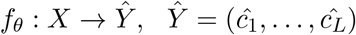

where each ĉ_*i*_ is chosen as a multi-class classification problem from the set of valid synonymous codons *V* (*a*_*i*_) ⊂ 𝒞. Next, we discuss each architectural sub-component of ppLM-CO illustrated in Fig. 1a.

**Figure 1.**
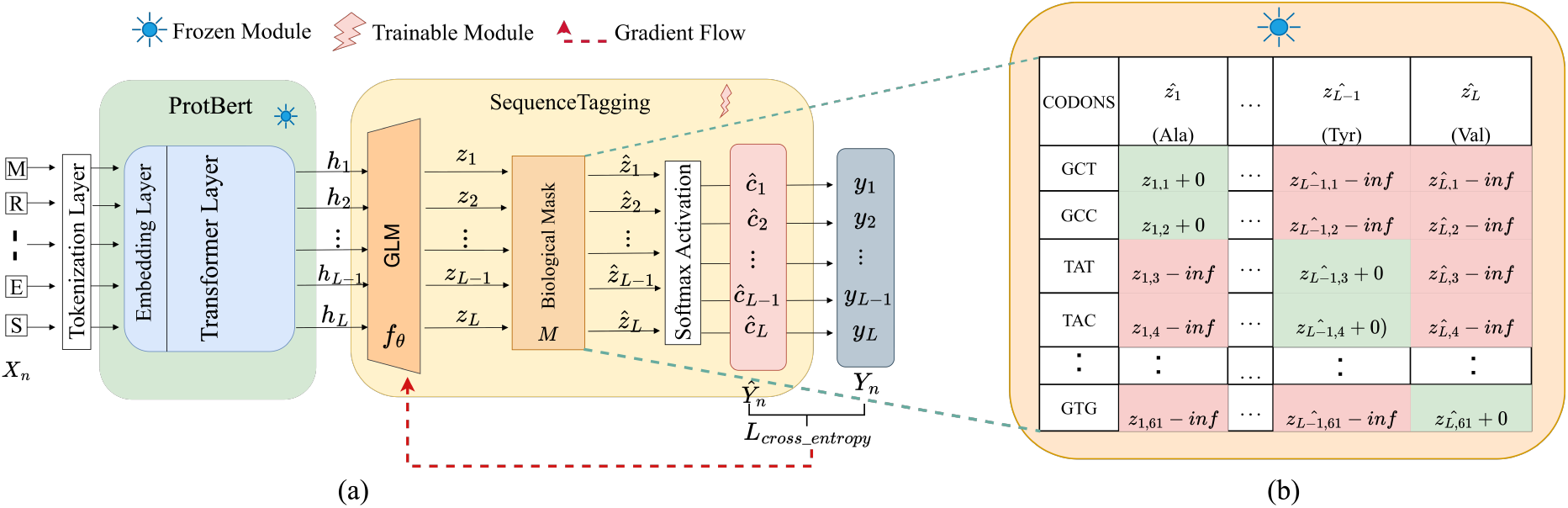
Illustration of the proposed ppLM-CO. Fig (a) illustrates different components in the ppLM-CO architecture. Fig (b) illustrates the biological mask where 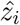 is the logits and (.) is its corresponding amino-acid. Cells shaded with green are synonymous codons for the corresponding amino-acid, while the red shaded cells belong to non-synonymous codons. For example, ‘Ala’ amino-acid synonymous codons as illustrated in fig.(b) are ‘GCT’, ‘GCC’.

#### Encoder

To capture the contextual relationship between input *a*_*i*_ in *X*_*n*_, in this work, we eliminate the need for training an encoder like RNN or transformers using a ppLM: ProtBert [17]. For each amino-acid *a*_*i*_, its contextual embedding from the encoder is obtained as:

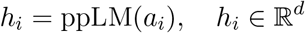

where d denotes the embedding dimension, here *d* = 1024 for ProtBert [17]. Recall we do not train the ppLM as an encoder and keep its parameters frozen. The resulting representation *H* = (*h*_1_, …, *h*_*L*_) captures the contextual relationship of amino-acids in the input protein sequence.

#### Codon Classifier (GLM Layer)

While the encoder extracts each amino-acid embedding, codon prediction in the sequence tagging task requires transforming this representation into codon-class probabilities. Therefore, we use a single linear layer classifier to map the aminoacids embedding to logits over 61 codons as follows:

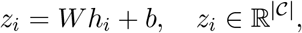

where, *W* ∈ ℝ^|*𝒞*|*×d*^ and *b* ∈ ℝ^|*𝒞*|^. This layer performs the multi-class classification conditioned on pre-trained amino-acid embedding (*h*_*i*_) coming from ppLM: ProtBert [17]. The GLM layer serves as the prediction head for codon selection and is the only trainable component in our work.

#### Biological Masking

Since the genetic code is degenerate, each amino-acid *a*_*i*_ corresponds only to a subset of valid codons: *V* (*a*_*i*_) ⊂.𝒞 Consequently, only a subset of logits *z*_*i*_ predicted by the classifier are biologically meaningful. To explicitly induce the genetic constraint, we perform a biological masking operation that restricts the model’s label space to valid synonymous codons (Fig. 1b). The mask matrix *M* ∈ ℝ^*L×*|*𝒞*|^ is defined as:

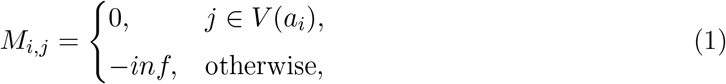

and applied over the classifier logits as: 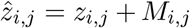. This biological mask acts as a genetic prior that enforces synonymous translation without predefining which codon is optimal. It does not simplify the learning task but constrains it to the biologically valid search space, allowing the model to focus on learning host-specific codon preferences rather than rediscovering the universal genetic code.

#### Prediction and Training Objective

After masking, codon probabilities (*c*_*i*_) are obtained through a softmax function [23]:

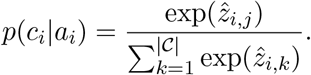

To train the model, we minimize the cross-entropy loss defined as: 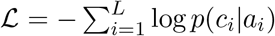 At inference, the optimal codon corresponding to *a*_*i*_ is chosen as: ĉ_*i*_ = arg max_*j*_ *p*(*c*_*i*_|*a*_*i*_).

### 2.2 Dataset

We curated datasets for three host species—Human, *E*.*coli*, and Chinese-Hamster—to train and evaluate species-specific instances of ppLM-CO. Each dataset provides paired amino-acid sequences and their corresponding coding sequences (CDSs), enabling supervised learning of host-dependent codon usage preferences.

For the Human dataset, we collected protein–CDS pairs from the UCSC (www.genome.ucsc.edu) and NCBI GenBank (www.ncbi.nlm.nih.gov/genbank). Following from previous work [14] we filter out sequences that are biologically invalid. We further retain sequences whose CDS length is between 250 and 2500 as long sequences can hinder training efficiency [16]. Second, we retain sequences with MFE ≤ −76 kcal/mol computed using ViennaRNA v2.6.4 [20]. This filtering ensures inclusion of structurally stable mRNAs so that ppLM-CO focuses on modeling codon usage patterns rather than transcript instability.

For *E*.*coli* and Chinese-Hamster we directly utilize the datasets from previous works [12, 14]. No MFE-based filtering was applied for these datasets as we primarily utilize them to validate cross-species generalization of ppLM-CO to capture distinct species-specific codon usage pattern. We provide each of the three datasets here:https://github.com/shashankcuber/Pre-trained-PPLM-Codon-Optimization.

## 3 Results

We comprehensively evaluate ppLM-CO *in-silico* across species and for the mRNA vaccine design to assess its effectiveness, generalization, and biological validity. All evaluations are conducted using widely accepted computational proxies of biological performance, including CAI for translational efficiency and MFE for mRNA structural stability.

Our evaluation proceeds in four stages. First, we benchmark on Human dataset, by training and evaluating ppLM-CO for expression (CAI) and stability (MFE) against state-of-the-art codon optimization methods including CodonBERT [16] and LinearDesign [21]. Second, we assess cross-species generalization by training and testing ppLM-CO on *E. col*i and Chinese-Hamster datasets, demonstrating its adaptability to capture species-specific codon usage patterns. Third, we apply ppLM-CO to mRNA vaccine CDS design, evaluating its optimized CDSs for SARS-CoV-2 spike protein and VZV gE protein against existing industry-approved vaccine sequences from Moderna (mRNA1273) and Pfizer(BNT162b2), as well as with CodonBERT [16] and LinearDesign [21]. Finally, we compare the trainable parameter counts of ppLM-CO with prior deep learning approaches like CodonBERT [16] and Codon-Box [12] to highlight its training computational efficiency.

### 3.1 Codon Optimization in Human

In this section, we discuss ppLM-CO performance against the Wild-Type CDS-coming from the Human dataset test-set and state-of-the-art methods: CodonBERT [16] and LinearDesign [21]. Our proposed ppLM-CO achieves the highest mean CAI (0.98 ± 0.02), surpassing CodonBERT [16] and LinearDesign [21](Fig. 2a) and Wild-Type CDSs. For stability, it reached mean MFE of −282 ± 212 kcal/mol-second to LinearDesign [21], which explicitly optimizes structural energy (Fig. 2a). The joint scatter plot of CAI-MFE in Fig. 2b, illustrates ppLM-CO predictions concentration in the high-CAI and low-MFE region, reflecting high gains in expression while maintaining competitive stability to previous works.

**Figure 2.**
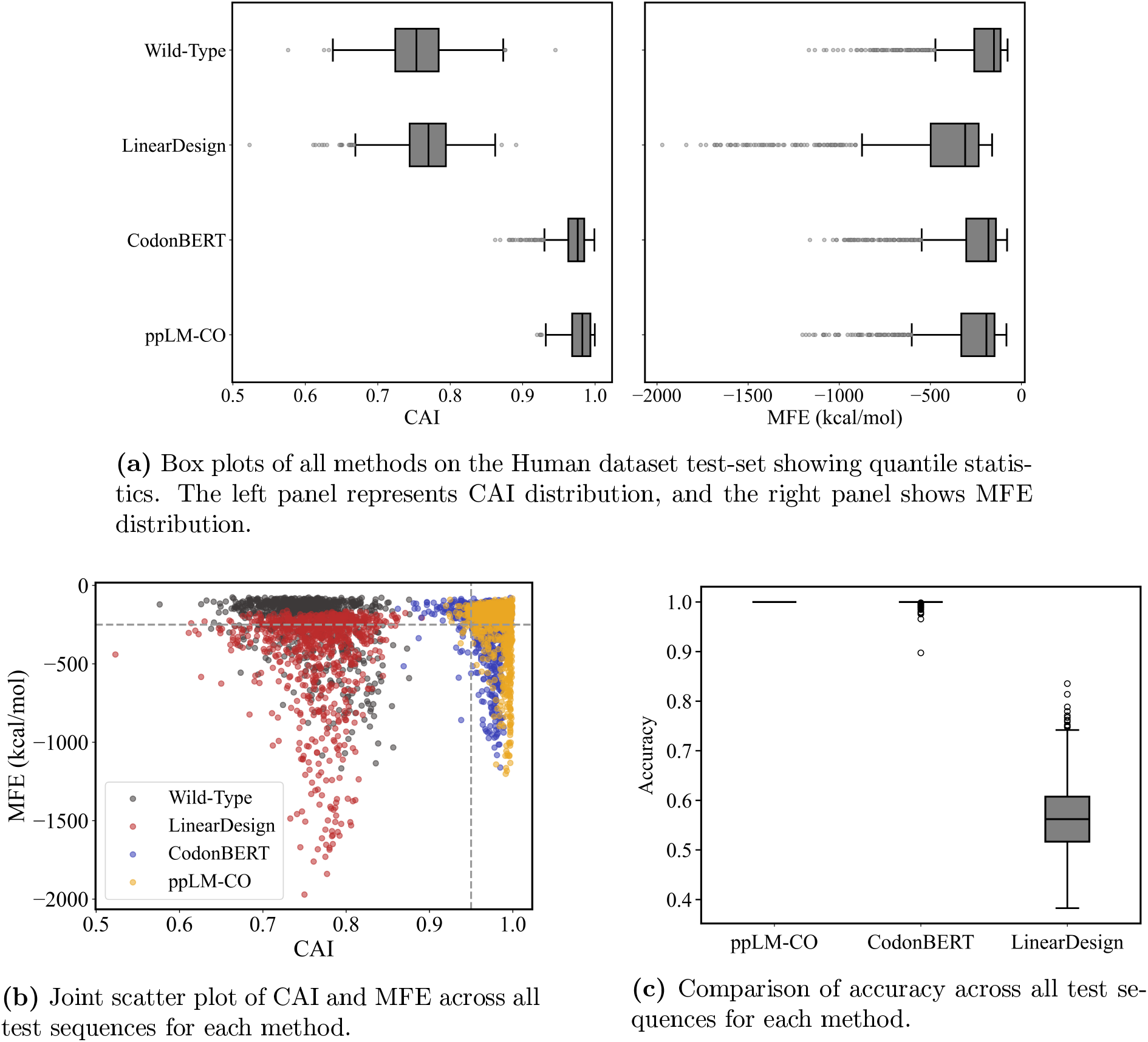
Evaluation of ppLM-CO against existing methods on the Human test-set. Subfigure (a) summarizes CAI and MFE distributions across methods using box-plots, Subfigure (b) illustrate CAI–MFE relationships highlighting expression–stability trade-offs. Subfigure (c) evaluates accuracy across methods.

We further evaluate the accuracy of ppLM-CO predictions on the Human test-set to quantify the proportion of non-synonymous codons in the optimized CDSs. We calculate the accuracy following from the work CodonBERT [16]. ppLM-CO achieves mean accuracy of 100% which is superior to previous approaches: CodonBERT (99.89%) [16], Codon-Box (52%) [12] and LinearDesign (56.51%) [21]. It indicates that all the CDS predictions from ppLM-CO were biologically valid and synonymous with target amino-acids, demonstrating the potential of proposed biological masking as a genetic prior during model training. Fig. 2c, illustrates the distribution of accuracy performance 820 test sequences, highlighting the perfect fidelity of ppLM-CO.

Our results in table 1 demonstrate that leveraging ppLM amino-acid embedding allows to capture human codon-usage patterns, yielding improvements with minimal training parameters.

**Table 1.** Results on the Human dataset test-set across different methods. The best and the second best metrics across methods are highlighted in **red** and **blue**, respectively.

| Method | CAI (mean ± std ↑) | MFE (mean ± std ↓, kcal/mol) |
| --- | --- | --- |
| Wild-Type (Baseline) | 0.75 ± 0.05 | -230 ± 188 |
| LinearDesign [21] | 0.77 ± 0.04 | -446 ± 337 |
| CodonBERT [16] | 0.97 ± 0.02 | -266 ± 202 |
| ppLM-CO(**Ours**) | 0.98 ± 0.02 | -282 ± 212 |

### 3.2 Cross-Species Generalization

To assess the generalization of ppLM-CO beyond human as a host species, we trained it independently on *E*.*coli* and Chinese-Hamster datasets and evaluated them against their respective Wild-Type CDSs in their respective test-sets. We do not compare with prior works in this section’s experiments as the goal is to validate the generalization of our proposed ppLM-CO to capture distinct host species codon usage pattern.

In E. coli, ppLM-CO improved the mean CAI from 0.67 to 0.97 and reduced the mean MFE by approximately 9 kcal/mol (Fig. 3a(1)). The CAI–MFE joint distribution shows optimized CDSs concentrated in the lower-right quadrant, representing higher expression and stability illustrated in Fig. 3a(2).

**Figure 3.**
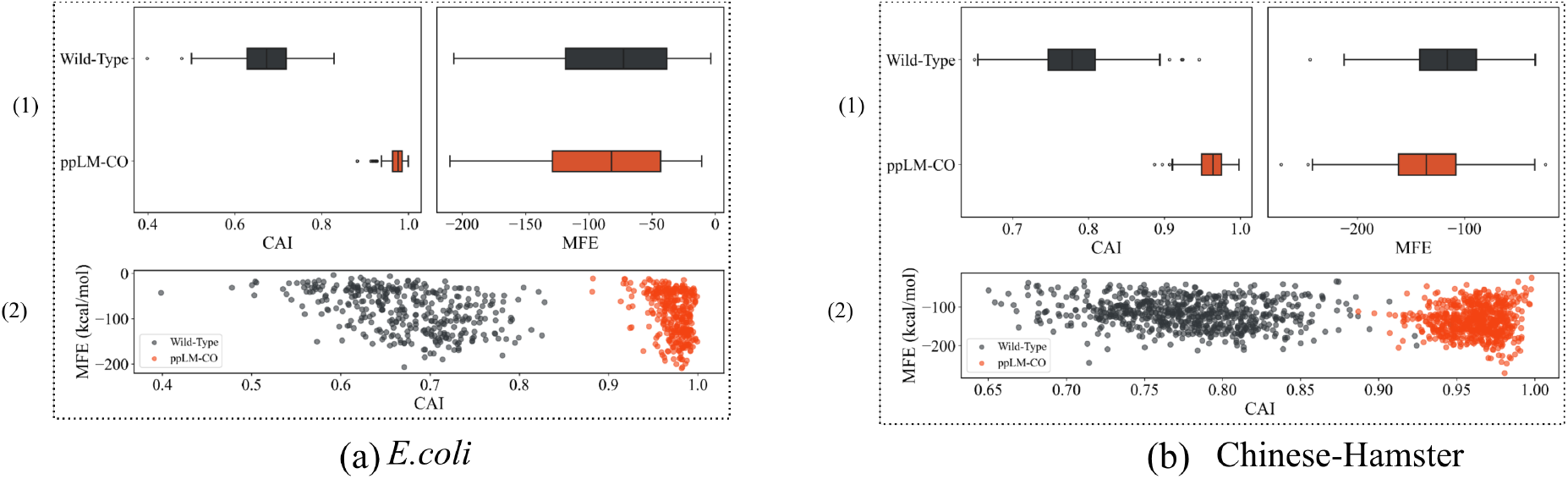
Cross-species generalization of ppLM-CO. (a) Performance of ppLM-CO compared to Wild-Type CDSs on the *E*.*coli* test-set. (b) Performance of ppLM-CO compared to Wild-Type CDSs on the Chinese-Hamster test-set. For each species, subfigure (1) contains the box plots of CAI and MFE, and subfigure (2) plots the joint CAI–MFE distribution of ppLM-CO optimized CDSs versus Wild-Type CDSs, illustrating consistent improvement in both expression and stability across species.

Similarly, in Chinese-Hamster, ppLM-CO increased mean CAI from 0.78 to 0.96 and lowered MFE by 19 kcal/mol compared to Wild-Type CDSs (Fig. 3b(1)). The resulting CDSs again cluster in the lower-right quadrant (Fig. 3b(2)), confirming consistent optimization across metrics. Overall, across all training and test datasets, every predicted CDS from ppLM-CO successfully reverse-translated to its corresponding input protein sequence. Additionally, the GC content of optimized sequences consistently remained within the biologically accepted range of 30 – 70% across all three species, reflecting balanced codon composition and avoiding extremes that can impair transcriptional or translational efficiency. Together, these findings demonstrate that ppLM-CO effectively captures host-specific codon usage patterns while maintaining biologically valid nucleotide distributions. The model generalizes robustly across species, offering a flexible and reliable framework for codon optimization that can be readily adapted to new host systems.

### 3.3 *In-Silico* mRNA Vaccine CDS Design

We next evaluate ppLM-CO for *in-silico* mRNA vaccine CDS design in humans, focusing on the SARS-CoV-2 spike protein and the VZV-gE protein. We utilize the ppLM-CO pre-trained on the Human dataset to generate CDS for the given two viral proteins. For VZV gE, industry vaccines from Pfizer and Moderna are unavailable for comparison.

As summarized in table 2, ppLM-CO achieved superior CAI across both antigens—0.99 for the SARS-CoV-2 spike and 0.99 for VZV gE—surpassing CodonBERT [16], LinearDesign [21], and even the optimized commercial vaccine CDSs from Pfizer (0.94) and Moderna (0.97). In terms of stability, ppLM-CO obtained MFE values of −1467 kcal/mol (SARS-CoV-2) and −744 kcal/mol (VZV), ranking second to LinearDesign [21].

**Table 2.** Comparison of methods for vaccine CDS design in humans as a host species. Results are shown for the SARS-CoV-2 spike protein^1^ and VZV gE protein^2^. CAI (Codon Adaptation Index, higher is better) measures codon usage bias, while MFE (Minimum Free Energy, lower is better) measures stability. The best values are in **red**, second-best in **blue**

| Method | SARS-CoV-2 |  | VZV |  |
| --- | --- | --- | --- | --- |
| | CAI (↑) | MFE (↓, kcal/mol) | CAI (↑) | MFE (↓, kcal/mol) |
| Wild-Type (Baseline) | 0.67 | -1111.5 | 0.66 | -485.7 |
| Pfizer (BNT162b2) | 0.94 | -1314.1 | — | — |
| Moderna (mRNA1273) | **0.97** | -1454.8 | — | — |
| CodonBERT [16] | 0.94 | -1341.1 | **0.97** | -706.3 |
| LinearDesign [21] | 0.72 | **-2477.7** | 0.77 | **-1249.10** |
| ppLM-CO (Ours) | **0.99** | **-1467.3** | **0.99** | **-743.7** |

These results demonstrate the potential of ppLM-CO as a general framework for mRNA therapeutics design. Despite its minimal parameter trainable module, it consistently generates biologically coherent, high-expression, and structurally stable CDSs across distinct viral proteins.

### 3.4 Computational and Parameter Efficiency of ppLM-CO

In the previous sections, we established the performance and generalization capabilities of our ppLM-CO across species and vaccine design. Unlike prior end-to-end deep models that retrain millions of parameters, ppLM-CO leverages a frozen pretrained protein language model—letting the model’s contextual amino-acid representations perform the heavy lifting—while fine-tuning only a lightweight classifier.

As shown in Table 3, ppLM-CO trains on just 62K parameters, which is 92% and 99% reduction in trainable parameter count compared to Codon-Box [12] and CodonBERT [16] respectively. Despite this large reduction in trainable parameters, it consistently outperforms previous methods. This efficiency makes ppLM-CO a practical, biologically grounded, and computationally lean framework for codon optimization.

**Table 3.** Training parameter count comparison with existing deep learning methods. The best and second-best values are highlighted in red and blue.

| Method | #Trainable Parameters (↓) |
| --- | --- |
| CodonBERT [16] | 7.1M |
| Codon-Box [12] | 829K |
| ppLM-CO (**Ours**) | 62K |

## 4 Conclusion

In this study, we present ppLM-CO, a lightweight training framework for codon optimization that leverages a pLM to capture host-specific translation biases. Specifically, ppLM-CO integrates pretrained protein language models (pLMs) to provide contextual amino acid embeddings and employs a biologically constrained classifier for synonymous codon prediction. *In-silico* performance across Human, bacterial (E. coli), and Chinese-Hamster datasets demonstrate superior expression (CAI) and competitive stability (MFE) while reducing trainable parameters by over 92% – 99% to previous works used for comparisons in this work. As future work, a possible extension could be to explore ppLM-CO towards structure-aware optimization for improving the stability and *in-vitro* mRNA vaccine and therapeutic design.

## Acknowledgements

This work was supported by the Natural Sciences and Engineering Research Council of Canada.

## References

[1] Vitiello, A. & Ferrara, F. Brief review of the mrna vaccines covid-19. Inflammopharmacology 29, 645–649 (2021).

[2] Chaudhary, N., Weissman, D. & Whitehead, K. A. mrna vaccines for infectious diseases: principles, delivery and clinical translation. Nature reviews Drug discovery 20, 817–838 (2021).

[3] Parhiz, H., Atochina-Vasserman, E. N. & Weissman, D. mrna-based therapeutics: looking beyond covid-19 vaccines. The Lancet 403, 1192–1204 (2024).

[4] Zarghampoor, F., Azarpira, N., Khatami, S. R., Behzad-Behbahani, A. & Foroughmand, A. M. Improved translation efficiency of therapeutic mrna. Gene 707, 231–238 (2019).

[5] Zhu, J. Mammalian cell protein expression for biopharmaceutical production. Biotechnology advances 30, 1158–1170 (2012).

[6] Kudla, G., Lipinski, L., Caffin, F., Helwak, A. & Zylicz, M. High guanine and cytosine content increases mrna levels in mammalian cells. PLoS Biology 4, e180 (2006).

[7] Kim, S. C. et al. Modifications of mrna vaccine structural elements for improving mrna stability and translation efficiency. Molecular & cellular toxicology 1–8 (2022).

[8] Calvo, S. E., Pagliarini, D. J. & Mootha, V. K. Upstream open reading frames cause widespread reduction of protein expression and are polymorphic among humans. Proceedings of the National Academy of Sciences 106, 7507–7512 (2009).

[9] Paremskaia, A. I. et al. Codon-optimization in gene therapy: promises, prospects and challenges. Frontiers in Bioengineering and Biotechnology 12, 1371596 (2024).

[10] Vaswani, A. et al. Attention is all you need. Advances in neural information processing systems 30 (2017).

[11] Goshisht, M. K. Machine learning and deep learning in synthetic biology: Key architectures, applications, and challenges. ACS omega 9, 9921–9945 (2024).

[12] Fu, H. et al. Codon optimization with deep learning to enhance protein expression. Scientific reports 10, 17617 (2020).

[13] Jain, R., Jain, A., Mauro, E., LeShane, K. & Densmore, D. Icor: improving codon optimization with recurrent neural networks. BMC bioinformatics 24, 132 (2023).

[14] Goulet, D. R. et al. Codon optimization using a recurrent neural network. Journal of Computational Biology 30, 70–81 (2023).

[15] Gong, H. et al. Integrated mrna sequence optimization using deep learning. Briefings in Bioinformatics 24, bbad001 (2023).

[16] Ren, Z. et al. Codonbert: a bert-based architecture tailored for codon optimization using the cross-attention mechanism. Bioinformatics btae330 (2024).

[17] Elnaggar, A. et al. Prottrans: Toward understanding the language of life through self-supervised learning. IEEE transactions on pattern analysis and machine intelligence 44, 7112–7127 (2021).

[18] Nijkamp, E., Ruffolo, J. A., Weinstein, E. N., Naik, N. & Madani, A. Progen2: exploring the boundaries of protein language models. Cell systems 14, 968–978 (2023).

[19] Carbone, A., Zinovyev, A. & Képes, F. Codon adaptation index as a measure of dominating codon bias. Bioinformatics 19, 2005–2015 (2003).

[20] Lorenz, R. et al. Viennarna package 2.0. Algorithms for Molecular Biology 6, 1–14 (2011).

[21] Zhang, H. et al. Algorithm for optimized mrna design improves stability and immunogenicity. Nature 621, 396–403 (2023).

[22] Xia, X. Detailed dissection and critical evaluation of the pfizer/biontech and moderna mrna vaccines. Vaccines 9, 734 (2021).

[23] Goodfellow, I., Bengio, Y. & Courville, A. Deep learning, chapter 6.2. 2.3 softmax units for multinoulli output distributions (2016).

